# Fast and Tuning-free Nonlinear Data Embedding and Integration based on DCOL

**DOI:** 10.1101/2024.06.06.597744

**Authors:** Shengjie Liu, Tianwei Yu

## Abstract

The rapid progress of single-cell technology has facilitated faster and more cost-effective acquisition of diverse omics data, enabling biologists to unravel the intricacies of cell populations, disease states, and developmental lineages. Additionally, the advent of multimodal single-cell omics technologies has opened up new avenues for studying interactions within biological systems. However, the high-dimensional, noisy, and sparse nature of single-cell omics data poses significant analytical challenges. Therefore, dimension reduction (DR) techniques play a vital role in analyzing such data. While many DR methods have been developed, each has its limitations. For instance, linear methods like PCA struggle to capture the highly diverse and complex associations between cell types and states effectively. In response, nonlinear techniques have been introduced; however, they may face scalability issues in high-dimensional settings, be restricted to single omics data, or primarily focus on visualization rather than producing informative embeddings for downstream tasks. Here, we formally introduce DCOL (Dissimilarity based on Conditional Ordered List) correlation, a functional dependency measure for quantifying nonlinear relationships between variables. Based on this measure, we propose DCOL-PCA and DCOL-CCA, for dimension reduction and integration of single- and multi-omics data. In simulation studies, our methods outperformed eight other DR methods and four joint dimension reduction (jDR) methods, showcasing stable performance across various settings. It proved highly effective in extracting essential factors even in the most challenging scenarios. We also validated these methods on real datasets, with our method demonstrating its ability to detect intricate signals within and between omics data and generate lower-dimensional embeddings that preserve the essential information and latent structures in the data.

## Introduction

Emerging single-cell technologies, such as high-throughput sequencing and high-resolution mass spectrometry, are providing more precise and cost-effective single-cell data for gene expression, protein abundance, chromatin accessibility, and DNA methylation and more [19]. An exponentially increasing amount of molecular measures in single cells across various tissues are generated with faster speed and higher precision. This surge empowers researchers to identify cell types, discover novel cell sub-groups, discern cell states, and trace developmental lineages.

While this vast amount of data holds the potential for valuable insights, omics datasets are typically high-dimensional and collected from thousands of cells, posing challenges for processing modeling algorithms. Additionally, Despite the numerous features, single-cell datasets often demonstrate sparsity, owing to technical limitations and lower intrinsic dimensionality of biological systems. Therefore, dimension reduction (DR) techniques, aiming at transforming high-dimensional data into lower-dimensional representations while preserving essential structure and characteristics, are indispensable when dealing with single-cell omics data.

Research on data dimension reduction has a long history. Principal Component Analysis (PCA), as one of the oldest and most widely used techniques, can be traced back to 1901 [22]. Besides PCA, there are other linear dimension reduction techniques such as Independent Component Analysis (ICA) [17] that linearly transforms variables into independent components with minimal statistical dependencies. Another technique is Linear Discriminant Analysis (LDA) [32], a supervised approach that extracts features linearly while considering class information to ensure maximum class separability.

However, the dependencies between biological features can be complex and far from linear. Linear DR approaches may not adequately capture the complexity of diverse cell types, leading to insufficient representation of the data. Consequently, numerous nonlinear DR techniques have been proposed, including kernel-based methods like Kernel-PCA (KPCA) [26], neural network-based methods such as Deep Count Autoencoder (DCA) [11], as well as methods focused on preserving local properties like Locally Linear Embedding (LLE) [23] and Laplacian Eigenmaps (LEIM) [5], and methods aimed at preserving global features such as distance-preserving techniques like Metric Multidimensional Scaling (MDS) [18], and its two variations Sammon Mapping [24] and Isomap [31]. Additionally, nonlinear DR methods like t-distributed Stochastic Neighbor Embedding (t-SNE) [33] and Uniform Manifold Approximation and Projection (UMAP) [21], which preserve the manifold structure of the data, has shown good results for many applications and are popular in single-cell analysis. However, these methods still have limitations. For example, the performance of Kernel-PCA depends on the choice of kernel function and can be computationally expensive, especially for large datasets. Methods based on or utilizing distance measures such as LLE, MDS, Sammon, t-SNE, and UMAP may be susceptible to the curse of dimensionality. In high-dimensional spaces, distances between data points may lose geometric meaning, affecting distance metrics and potentially biasing or distorting results. Among these nonlinear DR methods, t-SNE and UMAP are the most commonly used techniques in single-cell data analysis, but they are primarily employed for visualizing high-dimensional data in low dimensions rather than providing meaningful lower embeddings for downstream analysis. Additionally, despite their ability to provide relatively good results, their performances are sensitive to hyperparameter settings.

Meanwhile, technological advances have enabled simultaneous profiling of multiple types of omics on the same set of single cells. An increasing number of single-cell multimodal technologies are being developed, including single-cell methylome and transcriptome sequencing (scM&T-seq) [2], single-cell analysis of genotype, expression, and methylation (sc-GEM) [9], cellular indexing of transcriptomes and epitopes by sequencing (CITE-seq) [28], single-nucleus chromatin accessibility and RNA expression sequencing (SNARE-seq [20] and SHARE-seq [20]), 10x Multiome platform, etc.

Different omics data types provide unique views of cellular functioning, and integrating these measurements is thereby expected to provide a more comprehensive view of the biological system. The integration of multiple omics datasets can be achieved through various integrative methodologies, which can be broadly categorized. Network-based methods, exemplified by techniques like Similarity Network Fusion (SNF) [34] and weighted nearest neighbor (WNN) [13], construct sample-similarity networks for each omics layers and further fuse the resulting networks. In contrast, model-based integration methods such as the Multimodal Deep Neural Network by integrating Multi-dimensional Data (MDNNMD) proposed by Sun et al. [30], conduct separate analyses on each dataset and aggregate the findings into a single final prediction.

Joint Dimension Reduction (jDR) approaches integrate multiple omics data and produce either shared or omics-specific lower latent representations. Methods like Integrative non-negative matrix factorization (intNMF) [8], iCluster [27], MOFA [4], and its sequel version MOFA+ [3] decompose each omics dataset into the product of a factor matrix shared across all omics, alongside omics-specific weight matrices. Joint Singular Value Decomposition (jSVD) [25] simultaneously performs singular value decomposition on all omics datasets with a common cluster matrix *U*. Additionally, Canonical Correlation Analysis (CCA) [16] and its nonlinear extension kernel-CCA [1], as well as Co-Inertia Analysis (CIA) [10], infer omics-specific factors rather than a common sub-space, by maximizing measures of agreement between embeddings such as correlation or co-inertia. Furthermore, a number of neural network-based methods are designed to nonlinearly integrate and reduce dimensions of multimodal data. For instance, the Single-cell multimodal variational autoencoder (scMVAE) [36] integrates scRNAseq and scATAC-seq data using a variational autoencoder that encodes multi-omics data through a combination of three distinct methods. Aside from deep learning methods, which utilize complicated nonlinear mappings to integrate multi-omics data with limited interpretability, most joint dimension reduction (jDR) methods typically model associations between omics in a linear manner. Consequently, they lack the ability to capture nonlinear effects and interactions between different biological layers.

In this study, we introduce DCOL correlation, a general association measure designed to quantify functional relationships between two random variables. When two random variables are linearly related, their DCOL correlation essentially equals their absolute correlation value. When the two random variables have other dependencies that cannot be captured by correlation alone, but one variable can be expressed as a continuous function of the other variable, DCOL correlation can still detect such nonlinear signals. Based on this measure, we propose DCOL-PCA and DCOL-CCA, where we extended PCA for dimensionality reduction of single-omics data and CCA for integration and joint dimensionality reduction of paired-omics data.

Our method’s advantage lies in its ability to identify both linear and nonlinear structures within and between omics data, thus achieving effective integration and dimensionality reduction. Additionally, the association measure is based on the traditional statistical principle of assessing the percentage of variance explained, and DCOL-PCA and DCOL-CCA inherit the interpretability of PCA and CCA and are hyperparameter-free, eliminating the need to tune additional hyperparameters for optimal performance. The methods can be used not only for visualizing high-dimensional data but also for providing informative lower-dimensional embeddings for downstream tasks. We compared DCOL-PCA with eight other dimensionality reduction (DR) methods, and compared DCOL-CCA with four joint dimensionality reduction (jDR) methods on simulated data under various settings. Our methods achieved stable and superior performance across almost all scenarios. In real data analysis, DCOL-PCA and DCOL-CCA effectively capture intricate inter- and intra-relationships among omics data, yielding insightful lower-dimensional embeddings even in noisy datasets where other methods have failed to reduce dimensionality properly.

## Methods

### Dissimilarity based on Conditional Ordered List

We first introduced the dissimilarity based on conditional ordered list (DCOL) between a pair of random variables in [35]. Considering two random variables, *X* and *Y*, along with their respective data points 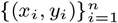, we organize these data points by arranging them in ascending order based on the values of *x*, resulting in: 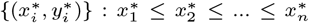. The dissimilarity between the variables *X* and *Y* is then calculated as:

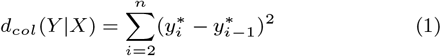

It is clear that when the spread of *Y* |*X* is small, the dissimilarity is small.

### DCOL Correlation

In this subsection, we derive the DCOL covariance between two random variables *X* and *Y*. Unlike conventional linear correlation, this measure accounts for both linear and nonlinear associations, providing a more holistic evaluation of the relationship between the two variables. Assume the relationship between Y and X is *Y* = *f* (*X*) + *ϵ*, where *f* () is a continuous function, and *ϵ* is the additive noise with mean 0 and variance *σ*^2^.

Utilizing DCOL, we can estimate *σ*^2^ without the need to estimate the specific functional form of *f* (). This is due to the relation:

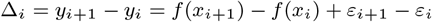

For sufficiently large sample sizes and assuming *X* is continuously distributed within a finite range:

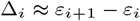

Furthermore, as the sample size n approaches infinity:

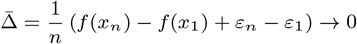

Under the assumption that *f* ()’s value and first derivative are finite everywhere, it can be demonstrated that:

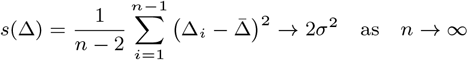

Thus, employing {Δ_*i*_}_*i*=1,…,*n*−1_, we can estimate the variance of *ϵ* as:

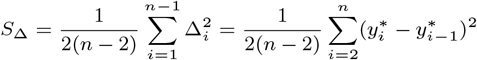

This *S*_Δ_ value serves as an approximation of *σ*^2^. Denoting the DCOL between *X* and *Y* as *d*_col_(*Y* |*X*), it can be expressed as:

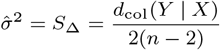

In regression analysis, by analyzing the source of variability, we have the following relationship:

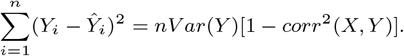

Let *See* = (*Y*_*i*_ − *Ŷi*)^2^ and 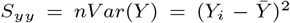, this equation can be equivalently expressed as:

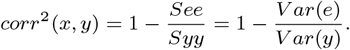

Using *S*_Δ_ as an estimation of the unexplained variability in situations where *Y* and *X* exhibit a relationship in the form of *Y* = *f* (*X*) + *ϵ*, we extend the previous outcome and define the DCOL-correlation score by substituting *V ar*(*e*) with *S*_Δ_:

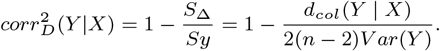

The DCOL-correlation score captures both linear and nonlinear relationships between two random variables. In practice, when *Y* and *X* are independent, DCOL-correlation follows a normal distribution with mean zero. Hence the DCOL-correlation score may sometimes fall below 0, indicating a lack of discernible functional association between the two random variables. In such cases, we set the DCOL-correlation score to zero, thereby ensuring it within the range of [0,1].

### DCOL-PCA

As the most widely used dimension reduction method, PCA reduces the dimensionality of data by constructing linear combinations of the original variables, aiming to capture and preserve the maximum variance. The basic idea of PCA is to find the first principal component that represents the most variation within the dataset. Subsequently, it seeks additional principal components that are uncorrelated with the previously identified components while accounting for the subsequent highest variance in the data. This process iterates until the new component becomes nearly insignificant or reaches a user-defined threshold.

One limitation of PCA is its focus on linear correlations, constraining its capability to capture nonlinear relationships among features. To address this constraint, we propose DCOL-PCA as an effective method for dimension reduction, particularly suited for scenarios where features present a mix of linear and nonlinear relationships.

Consider a data matrix *X* of size *n × p*, where *n* rows represent different samples, and *p* columns denote distinct features. Hereafter, we assume that the columns of matrix *X* have been standardized to have zero mean and unit variance. Dcol-PCA aims to find an orthogonal matrix *V* ∈ *R*^*d×p*^(*d << p*), which maps each *x*_*i*_ ∈ *R*^*p*^ to a lower dimensional space such that most relationships between features charecterized by the DCOL-correlation are retained.

Mathematically, it can be formulated as:

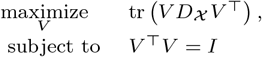

where *D*_*X*_ represents the DCOL-correlation matrix of *X*, with its (i,j)-th element defined as the larger value between corr_*D*_(**x**_*i*_|**x**_*j*_) and corr_*D*_(**x**_*j*_ |**x**_*i*_).

### DCOL-CCA

In addition to DCOL-PCA, we introduce another jDR method called DCOL-CCA, specifically designed for paired datasets. DCOL-CCA is an adaptable dimension reduction and multiomics integration approach which leverages the proposed DCOL-correlation measure and extends the Canonical Correlation Analysis (CCA) framework. It is capable of modeling general dependencies between paired data, thereby allowing for broader relationships beyond the linear constraints inherent in traditional CCA.

Let 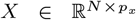 and 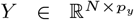 be two zero-mean and unit-variance data matrices with *N* instances and *p*_*x*_ and *p*_*y*_ features, respectively. DCOL-CCA aims to find two projection matrices 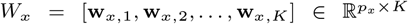 and 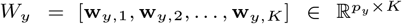, such that the DCOL-correlations between *XW*_*x*_ and *Y W*_*y*_ are maximized.

Take one vector 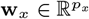 for *X* and one vector 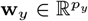 for *Y* as example. DCOL-CCA finds **w**_*x*_ and **w**_*y*_ by solving:

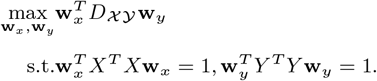

In omics data, the feature dimension is often high (*N << p*_*x*_ and *N << p*_*y*_), resulting in singular covariance matrices *X*^*T*^ *X* and *Y* ^*T*^ *Y*. As a consequence, the optimization problem becomes underdetermined. To remedy this problem, we introduce regularizations by adding two diagonal matrices:

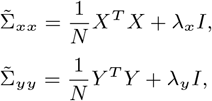

where *λ*_*x*_ and *λ*_*y*_ are non-negative regularization coefficients.

One way of computing *W*_*x*_ and *W*_*y*_ is to solve the following generalized eigenvalue decomposition problem:

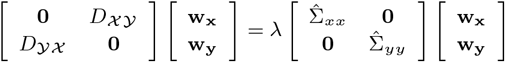

Here, the matrices 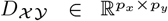 and 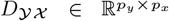 indicate the DCOL-correlation between *X* and *Y*, and *Y* and *X*, respectively. The (i,j)-th element of *D* _*𝒳 𝒴*_ is calculated as *max*(corr_*D*_(**x**_*i*_|**y**_*j*_), corr_*D*_(**y**_*i*_|**x**_*j*_)). The elements of *D* _*𝒴 𝒳*_ are similarly defined. The set of vectors 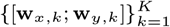 are then the *K* leading generalized eigenvectors.

### Simulation Study

#### Single-omics Data

For single-omics case, we generated data consisting of n samples ∈ ℝ^*p*^, categorized into k groups based on their features, implying an effective dimension of *k << p*. Within each group of size m, the features are nonlinearly associated. These nonlinear associations are randomly chosen from a set of six functions: (1) *f* (*x*) = cos(2*x*), (2) *f* (*x*) = *x*^2^, (3) *f* (*x*) = |*x*|, (4) *f* (*x*) = *x*^3^, (5) *f* (*x*) = 𝟙(−0.5 ≤ *x* ≤ 0.5), (6) *f* (*x*) = *atan*(4*x*).

The given pseudocode illustrates the simulation procedure. We conducted simulations across 27 scenarios, where three parameters were varied: the sample size *n* chosen from {500, 1000, 2000}, the number of features *p* selected from {500, 1000, 2000}, and the group size *k* picked from {5, 10, 15}. We utilized the Adjusted Rand Index (ARI) as the performance metric, quantifying the correspondence between the true labels and the clustering assignments. Our DCOL-PCA was compared against eight other dimension reduction methods: PCA, KPCA, LEIM, LLE, MDS, Sammon, t-SNE, and UMAP. For each simulation configuration, we conducted 10 experiments and computed the average ARI values.

##### Algorithm 1

Single-omics Data Simulation

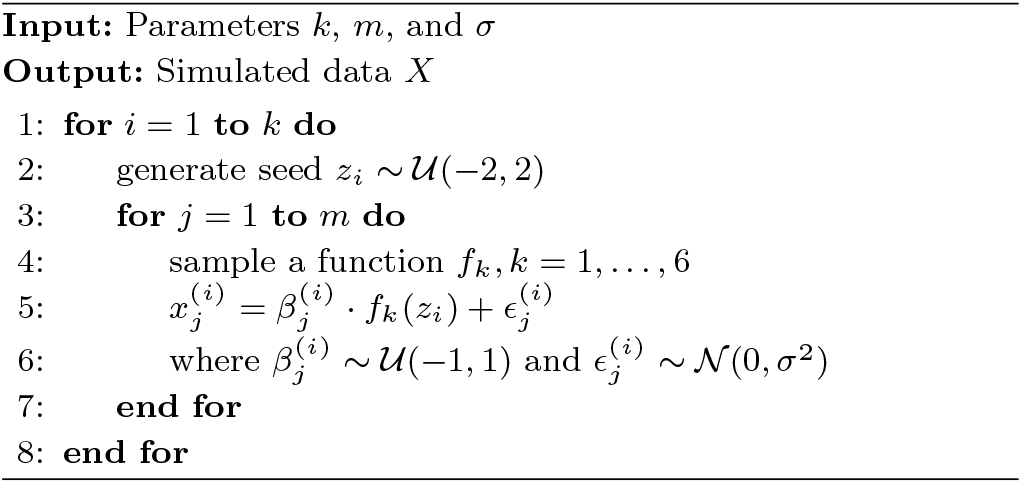

#### Paired-omics Data

In our study of paired-omics data, we generated two random matrices, denoted as *X* and *Y*, where the first *kx* variables of *X* are pair-wise associated with the first *kx × ky* variables of *Y*. The relationships between *X* and *Y* are randomly composed by the following six functions: (1) *f* (*x*) = *αx*, where *α* ∼ *𝒰* (−2, 2), (2) *f* (*x*) = *x*^2^, (3) *f* (*x*) = |*x*|, (4) *f* (*x*) = *sin*((*x* − 0.5) ∗ *π*), (5) *f* (*x*) = 𝟙(−0.5 ≤ *x* ≤ 0.5), (6) *f* (*x*) = *cos*^4^(0.75*x*). We describe the simulation procedure in the following pseudocode:

##### Algorithm 2

Paired-omics Data Generation

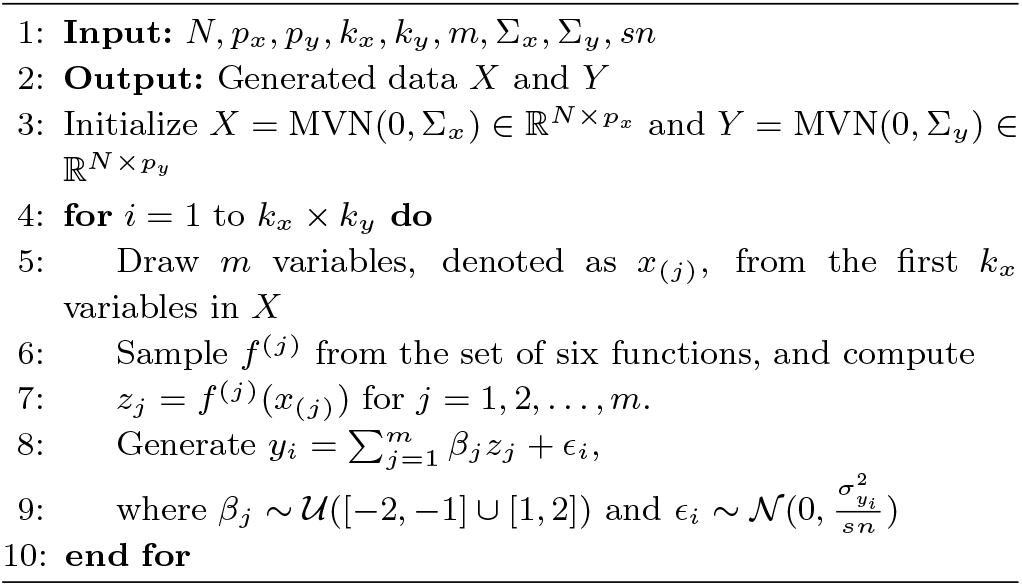

Here, ∑_*x*_ and ∑_*y*_ are matrices whose diagonal elements are 1 and off-diagonal elements take the value 0.1. We fix *p*_*x*_ and *p*_*y*_ at 1000 and set *m* = 4 and *sn* = 3.16. We analyzed the simulated data using five methods: DCOL-CCA, CCA, iCluster, jSVD, and MOFA+. To evaluate the methods, we first obtained the variable importance ranking generated by each method, and based on that, we computed the area under the ROC curve (AUC score) and the area under the precision-recall curve (PR-AUC score). A number of scenarios, specified by the combinations of *k*_*x*_, *k*_*y*_, and *N*, were conducted. In each scenario, the simulation was repeated 10 times.

### Simulation Results

#### Single-omics Data

The simulation results for the case of single-omics data are shown in Fig 1. We calculated the Adjusted Rand Index (ARI) scores based on the first five dimensions of the reduced space generated by individual dimension reduction methods. For methods like KPCA, t-SNE and UMAP, which involve tuning parameters, we assess two distinct sets of tuning parameters for each method, labeled as kPCA1 and kPCA2, etc.

**Fig. 1:**
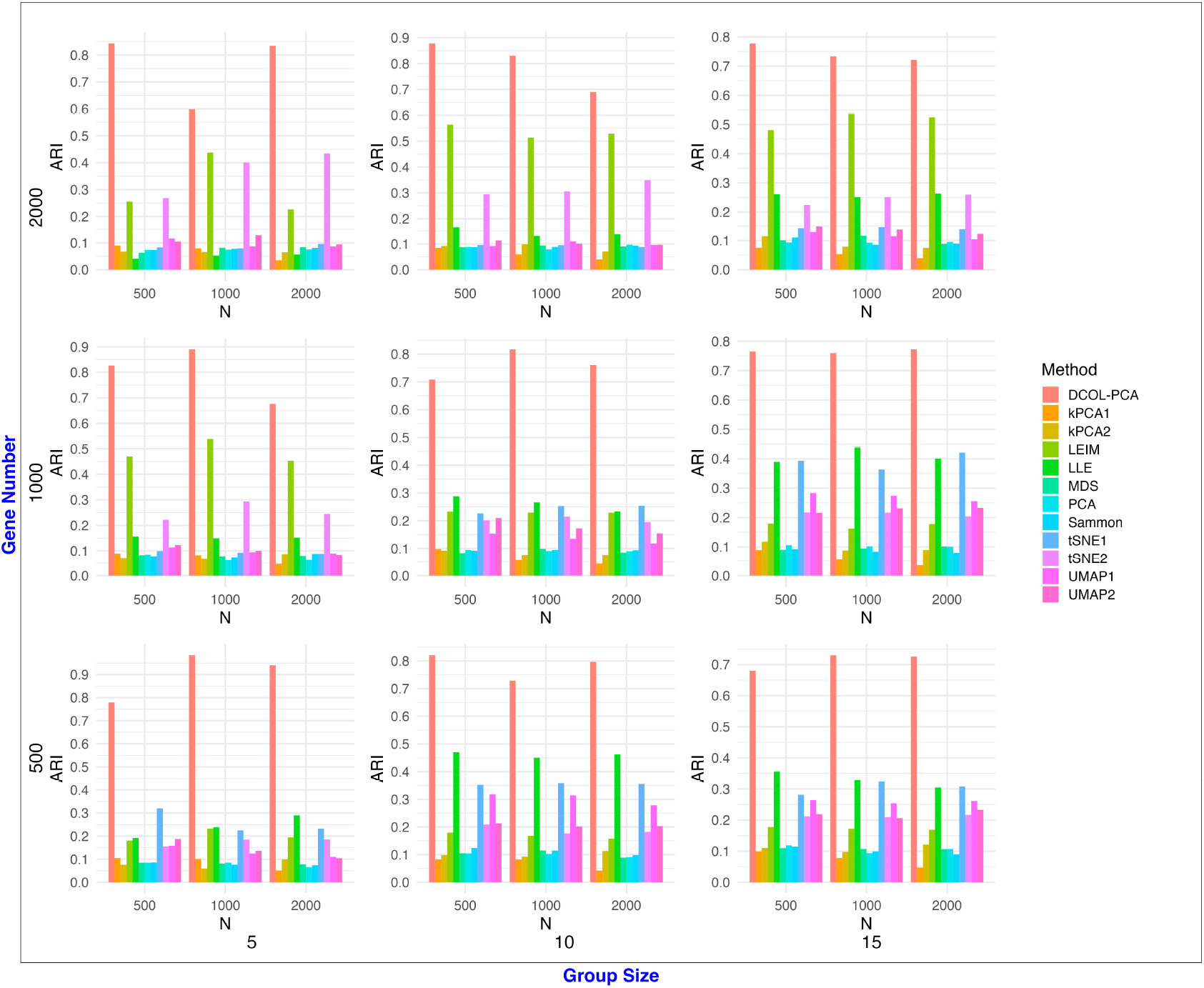
Single-omics simulation results. We conducted various simulations with different combinations of sample size (n) chosen from {500, 1000, 2000}, the number of features (p) selected from {500, 1000, 2000}, and the group size (k) picked from {5, 10, 15}. Adjusted Rand Index (ARI) scores were calculated based on the first five dimensions of the reduced space generated by each DR method.

Across all parameter settings, DCOL-PCA consistently outperforms the other eight methods, with the lowest ARI score around 0.7 and the highest reaching 0.9343. Notably, its performance remains relatively stable regardless of changes in the sample size, the number of groups and the number of genes, indicating robust performance in various conditions.

Among the other eight methods, LEIM, LLE, t-SNE, and UMAP demonstrated relatively good performance in different scenarios. Specifically, LEIM stood out when there was a large number of genes (= 2000) and an above-moderate number of feature groups, or when the gene count was moderate (= 1000) and the number of feature groups was relatively low. In these cases, LEIM achieved remarkable results and obtained the second-highest ARI score. Conversely, when confronted with fewer genes and a greater number of feature groups, LLE and tSNE outperformed the other six methods. In scenarios with a moderate gene count and a large group number, these two methods exhibited comparable performance, achieving an ARI score around 0.4. The remaining methods, except for UMAP which was able to identify some nonliner signals in certain settings, cannot capture inter-group nonlinear relationships and hence failed to achieve effective dimension reduction.

To facilitate a more intuitive comparison of the effectiveness of various dimension reduction algorithms, we opted for a scenario with smallest number of genes and samples, alongside a moderate number of feature groups. Specifically, this scenario including 500 genes, 500 samples and 10 feature groups, which is a non-trivial case. We visualized two factors of the embedded space produced by each method, with different colors denoting distinct groups. For KPCA, t-SNE, and UMAP, which employed different tuning parameters, we presented the results that showed higher ARI scores. The outcomes are illustrated in Fig 2.

**Fig. 2:**
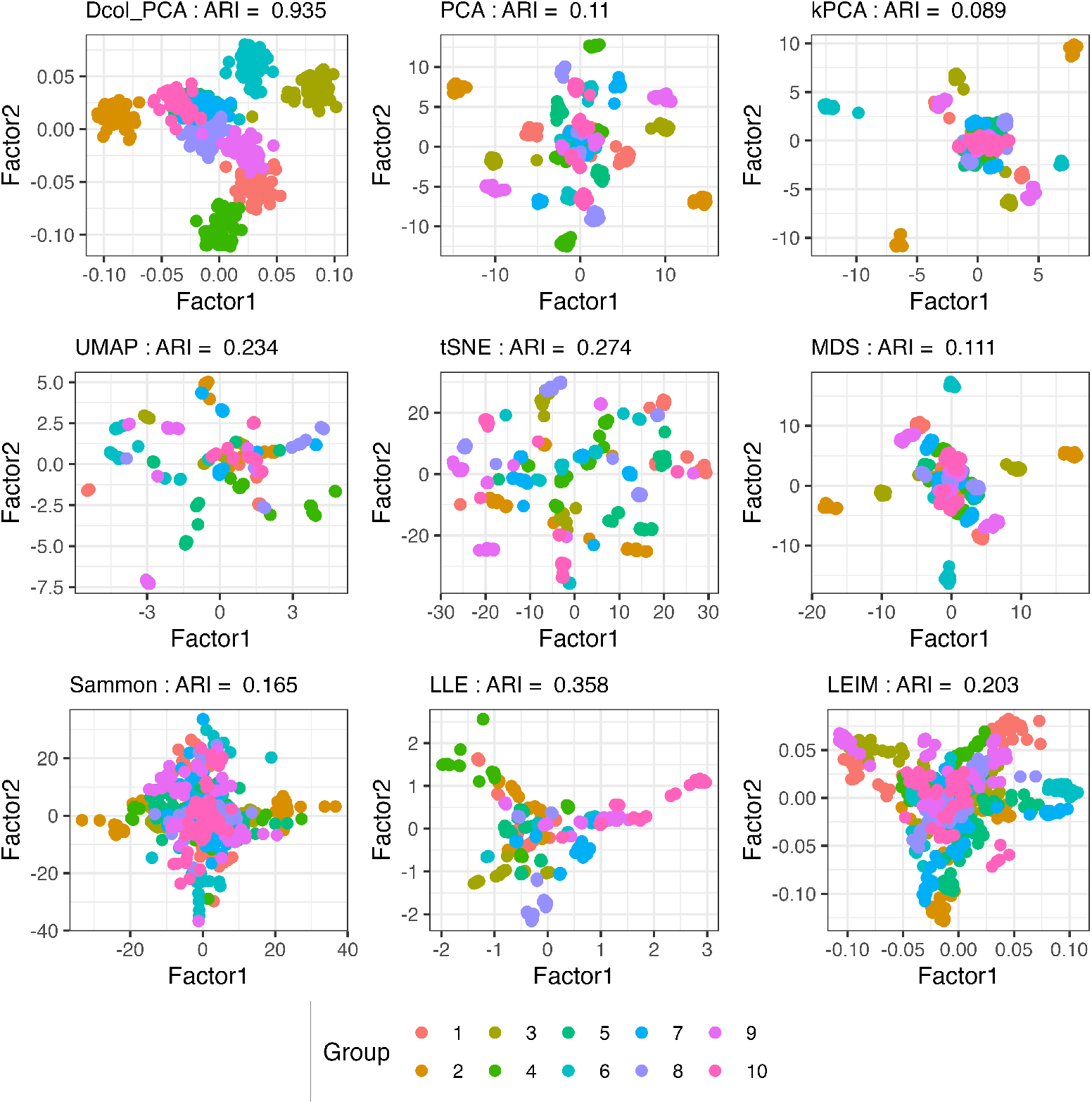
Comparison of dimension reduction methods. Two factors of the embedded space produced by each DR method are depicted, with distinct group colors. This scenario represents the smallest number of genes and samples, alongside a moderate number of feature groups (500 genes, 500 samples, and 10 feature groups)

**Fig. 3:**
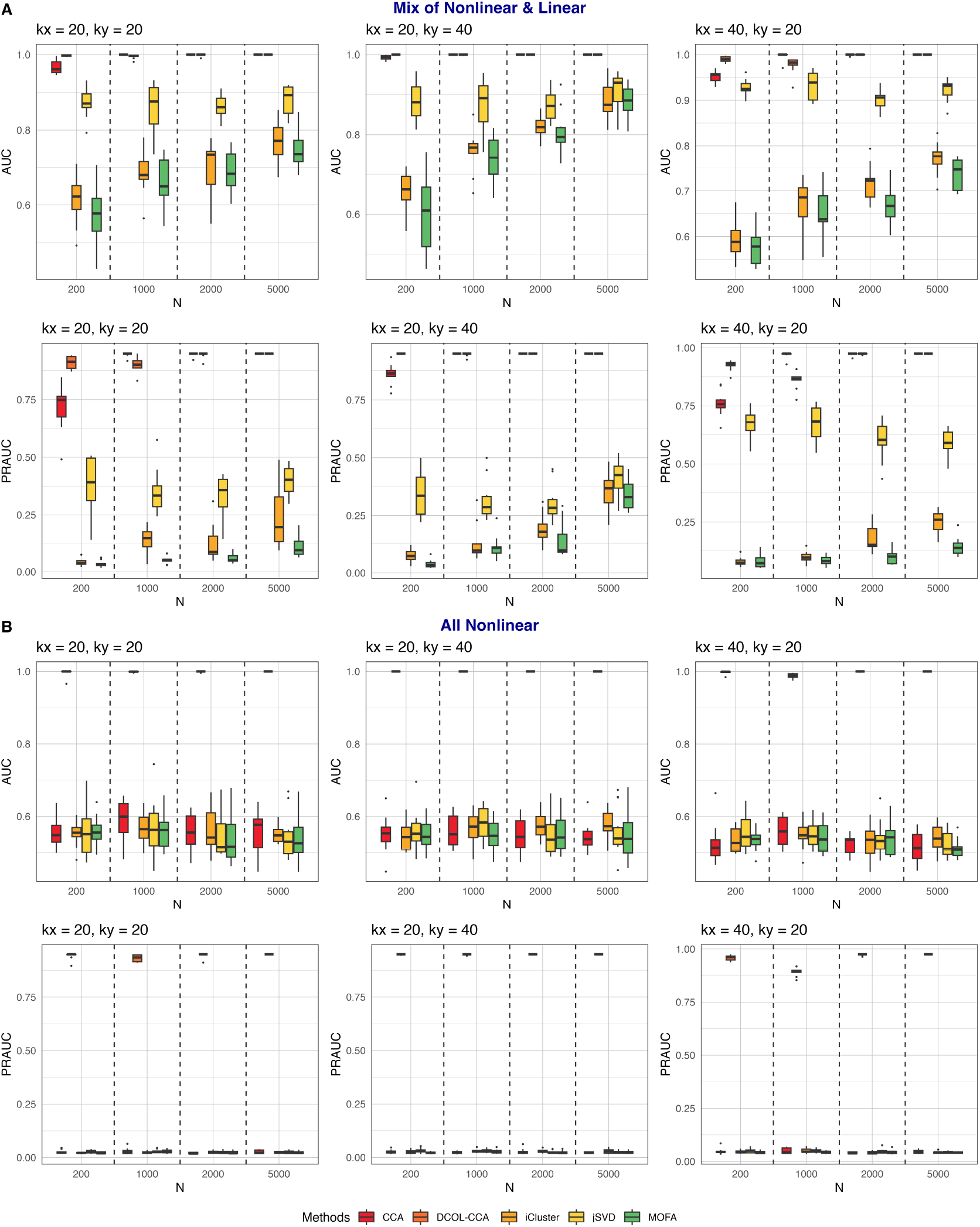
Simulation results of paired-omics. The PR-AUC metric was used to evaluate each method’s effectiveness in identifying the true contributing variables. Panel (a) features purely nonlinear relationships, while panel (b) shows a mix of nonlinear and linear relationships. X-axis: sample size; Y-axis: PR-AUC values.

Despite the slight overlap among groups 5, 6, and 7, we can distinctly observe different colors uniformly marked on separate clusters in the two factors generated by DCOL-PCA. This demonstrates our method’s agility in identifying general inter-group dependencies, allowing for effective dimensionality reduction that maximally preserves the group information in the reduced space. However, the results from other methods either display numerous dispersed small clusters, far exceeding the number of actual groups, or conglomerate into a nearly indistinguishable large cluster. The colors representing the groups are also haphazardly scattered across various clusters. Among them, LLE performed relatively well with the second-highest ARI score. In its outcome, although there were numerous small clusters, clusters sharing the same color stayed quite close. Nevertheless, the within-group relationship identified by LLE remained limited, impeding its ability to restore group information in the reduced space and leading to unsatisfactory performance. Similar patterns were observed across the results of other cases.

#### Paired-omics Data

The simulation results for the paired-omics data are shown in Fig.3. We devised two scenarios: (1) Mix of Nonlinear and Linear: where a small fraction of the initial *kx × ky* Y variables are non-linearly linked with the first *kx* X variables, while the rest demonstrate primarily linear correlations; and (2) All nonlinear: where the relationships between the first *kx* X variables and the first *kx × ky* Y variables are purely nonlinear.

When the relationships between X and Y are mixed (scenario 1), both CCA and DCOL-CCA exhibited the most optimal performance among all methods. DCOL-CCA demonstrated remarkable stability across various simulation settings, consistently outperforming CCA in nearly all cases. DCOL-CCA achieved AUC scores approaching or equaling 1 in almost all cases, while maintaining PRAUC scores consistently above 0.9, with the exception of a single case (when *kx* = 40, *ky* = 20, and *n* = 200). Among the remaining methods, jSVD showed the best performance, with all AUC values exceeding 0.85. However, its PRAUC scores lagged far behind the first two methods, particularly when *kx* = 20. When more X variables are associated with Y variables (*kx* = 40), it obtains better performance. Even when the relationships between X and Y are predominantly linear, iCluster and MOFA struggled to effectively identify X variables related to Y. Nevertheless, their performance displayed an upward trend with an increase in sample size.

When the relationship between *X* and *Y* is purely nonlinear, the performance of CCA, iCluster, jSVD, and MOFA dropped significantly. Despite their AUC scores ranging between 0.5 and 0.6, most of the simulated *X* variables do not actually contribute to the relationship with *Y*. Consequently, these methods may gain some AUC scores by erroneously identifying more contributing *X* variables than actually present. In turn, their PRAUC scores, all below 0.05, indicate their inability to distinguish the truly contributing *X* variables from the rest. Increasing the sample size did not lead to any improvement in their performances. In contrast, DCOL-CCA exhibited superior and reliable performance across all settings, with AUC scores approaching 1 and PRAUC scores consistently above 0.95. Even with a small sample size (=200), DCOL-CCA could still detect the relationship between *X* and *Y* variables and effectively pick out the 20 or 40 contributing *X* variables from among 1000 choices.

### Real Datasets

#### Single-omics Data

We compared DCOL-PCA against eight other methods using two real datasets with groud truth cell type labels: Segerstolpe and Bian, both downloaded from Cell Blast [7] (https://cblast.gao-lab.org/download). To assess the performance of various methods, we applied k-means to the initial five dimensions of the embedding produced by each method and obtained cell clusters. We adopted two metrics, Adjusted Rand Index (ARI) and Normalized Mutual Information (NMI), to quantify the extent to which the generated clusters align with the ground truth cell labels.

##### Segerstolpe dataset

The Segerstolpe dataset contains transcriptional profiles from individual pancreatic endocrine and exocrine cells of both healthy and type 2 diabetic donors. The dataset includes 1070 samples and 25453 genes, with samples classified into 12 subpopulations. We performed simple data preprocessing, which involved selecting the top 7000 high-variable genes and scaling the data to have unit variances and zero means.

Two factors of the lower-dimensional embeddings generated by the nine methods are displayed in Fig. 4A. Each point represents a cell line, and different colors indicate distinct cell populations. DCOL-PCA beats other methods with the highest ARI score (0.63) and NMI score (0.673). We observe seven consistently colored clusters from the result of DCOL-PCA. The clusters corresponding to cell populations alpha, gamma, and delta are isolated from the rest. Although the clusters of acinar, ductal, and beta stay close, they are separated from each other and uniformly colored. Notably, there are only five cell lines belonging to the epsilon population. However, DCOL-PCA still preserves the information of this tiny population and ensures they are close in the reduced space (indicated by the red circle in the subplot in the top-left corner of Fig. 4A). In the remaining methods, UMAP, t-SNE, and LEIM showed reasonably good performances, revealing discernible cell populations in their results. UMAP stood out with the highest ARI (0.509) and NMI (0.659) scores, while the ARI and NMI scores of t-SNE and LEIM were comparable, both around 0.4 and 0.6, respectively.

**Fig. 4:**
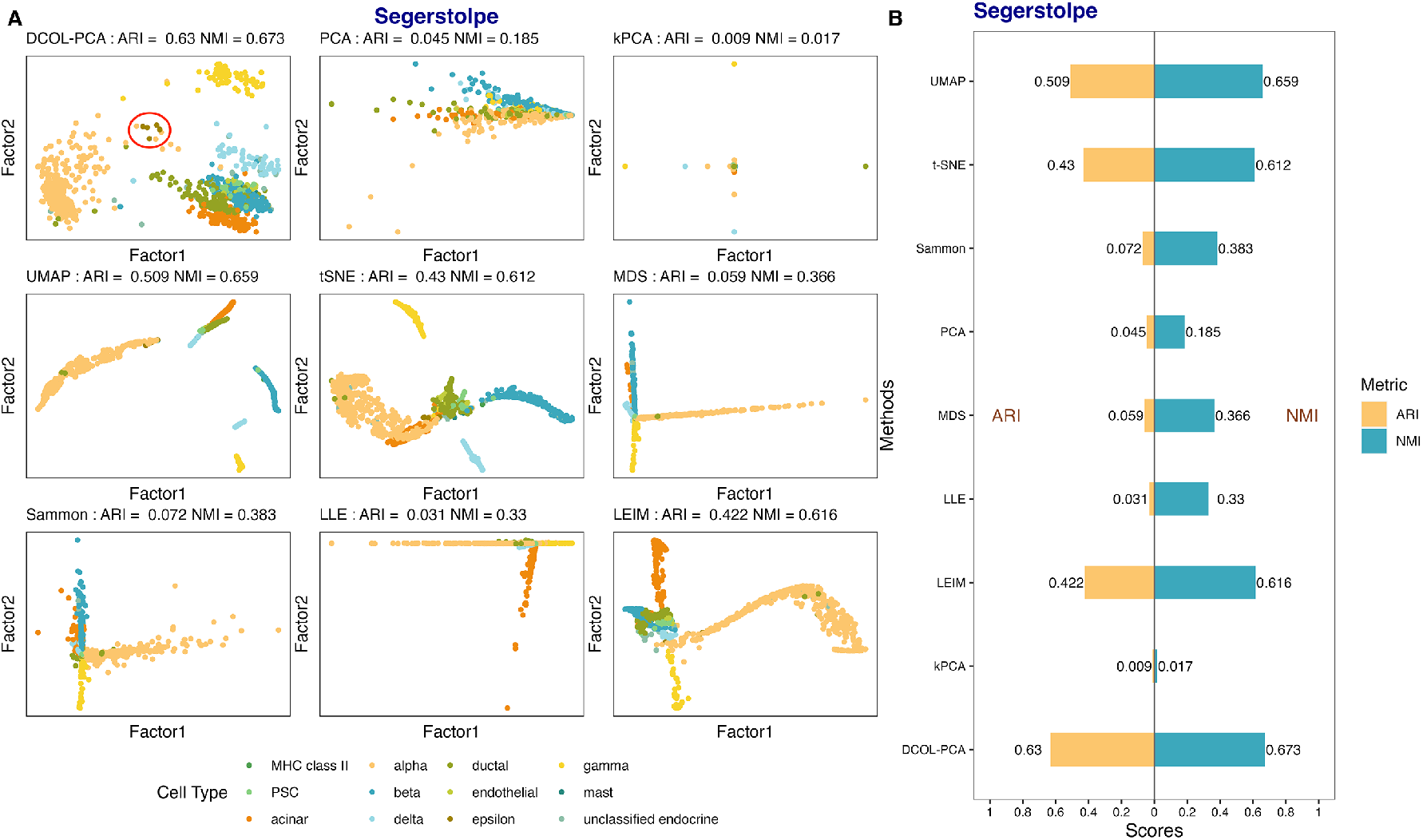
Results of DR methods on Segerstolpe dataset. (a) Scatterplots of two factors of the lower-dimensional embeddings are reported for each DR method. Each point represents a cell line and the colors represent distinct cell populations. (b) K-means clustering was applied to the initial five dimensions of each method’s embedding to obtain cell clusters. Performance was evaluated using the Adjusted Rand Index (ARI) and Normalized Mutual Information (NMI) to measure alignment with ground truth cell labels.

##### Bian dataset

The author of the Bian dataset sequenced CD45+ haematopoietic cells from human embryos at Carnegie stages 11 to 23 using single-cell RNA sequencing [6]. The dataset contains 1231 cell lines and 7371 genes. We followed the same preprocessing steps and evaluation procedure as the Segerstolpe dataset. The results of different DR methods are depicted in Fig. 5. This is a challenging dataset: ARI and NMI values for all methods are below 0.3. It can be observed that in the results of kPCA, UMAP, tSNE, LLE, and LEIM, colors representing cell populations are randomly distributed in the plots. In the results generated by PCA, MDS, and Sammon, the Mast cell (yellow) can be slightly distinguished. Nevertheless, even with the intricate relationships among features and within sub-populations, DCOL-PCA manages to discern signals and successfully identify various cell groups, including Myeloblast, Monocyte, Mac 1, Mac 2 and Mac 4.

**Fig. 5:**
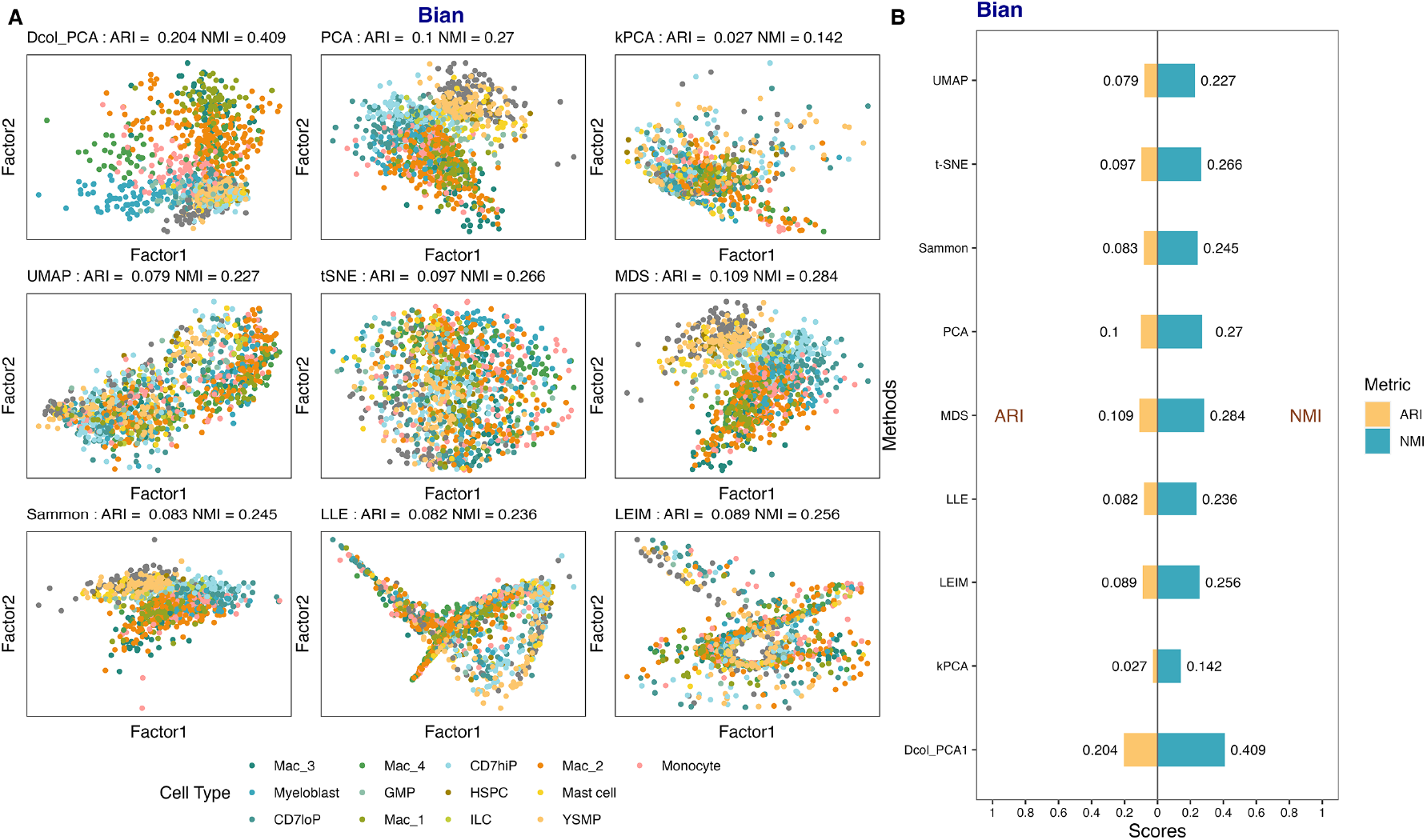
Results of DR methods on Bian dataset. (a) Scatterplots of two factors of the lower-dimensional embeddings for each DR method. (b) K-means clustering applied to initial five dimensions of each method’s embedding to obtain cell clusters. Performance assessed using Adjusted Rand Index (ARI) and Normalized Mutual Information (NMI) to quantify alignment with ground truth cell labels.

#### Paired-omics Data

We analyzed the an openly accessible dataset that jointly profiles m-RNA levels and DNA accessibility in single human peripheral blood mononuclear cells (PBMC), produced by 10x [**?**]. We computed per-cell quality control (QC) metrics using the DNA accessibility assay, including the strength of the nucleosome banding pattern and transcriptional start site (TSS) enrichment score. Cells failing to meet quality standards based on these metrics were filtered out using Signac [29], resulting in a dataset consisting of 10,412 cells. To identify a more accurate set of peaks, we called peaks using MACS2 [12] with the “CallPeaks” function. The dimensionality of resulting peak assay was reduced using latent semantic indexing (LSI) through Signac. Subsequently, we normalized the gene expression data using SCTransform and performed dimensionality reduction using PCA via Seurat [15].

We then annotated the cell types by transferring an annotated PBMC reference dataset from Hao et al. [14] using tools provided by Seurat. We applied CCA, DCOL-CCA, iCluster, jSVD and MOFA+ on the PBMC dataset.

Fig 6A shows the pair-wise plots of the first three embeddings returned by each method, with each point representing a cell line. Both CCA and DCOL-CCA effectively isolated pDC cells (depicted in light purple) from the remaining cell types. Furthermore, DCOL-CCA, jSVD, and MOFA+ successfully distinguished cDC2 and CD14 Mono cells. In addition, DCOL-CCA demonstrated exceptional performance in segregating various cell types, including B naive, B intermediate, CD16 Mono, NK, and MAIT cells, from the heterogeneous dataset.This suggests that DCOL-CCA adeptly integrated information from both modalities and achieved robust discrimination between cell groups. In contrast, the results obtained from other methods displayed greater overlap among data points, indicating a lower degree of cell type discrimination.

**Fig. 6:**
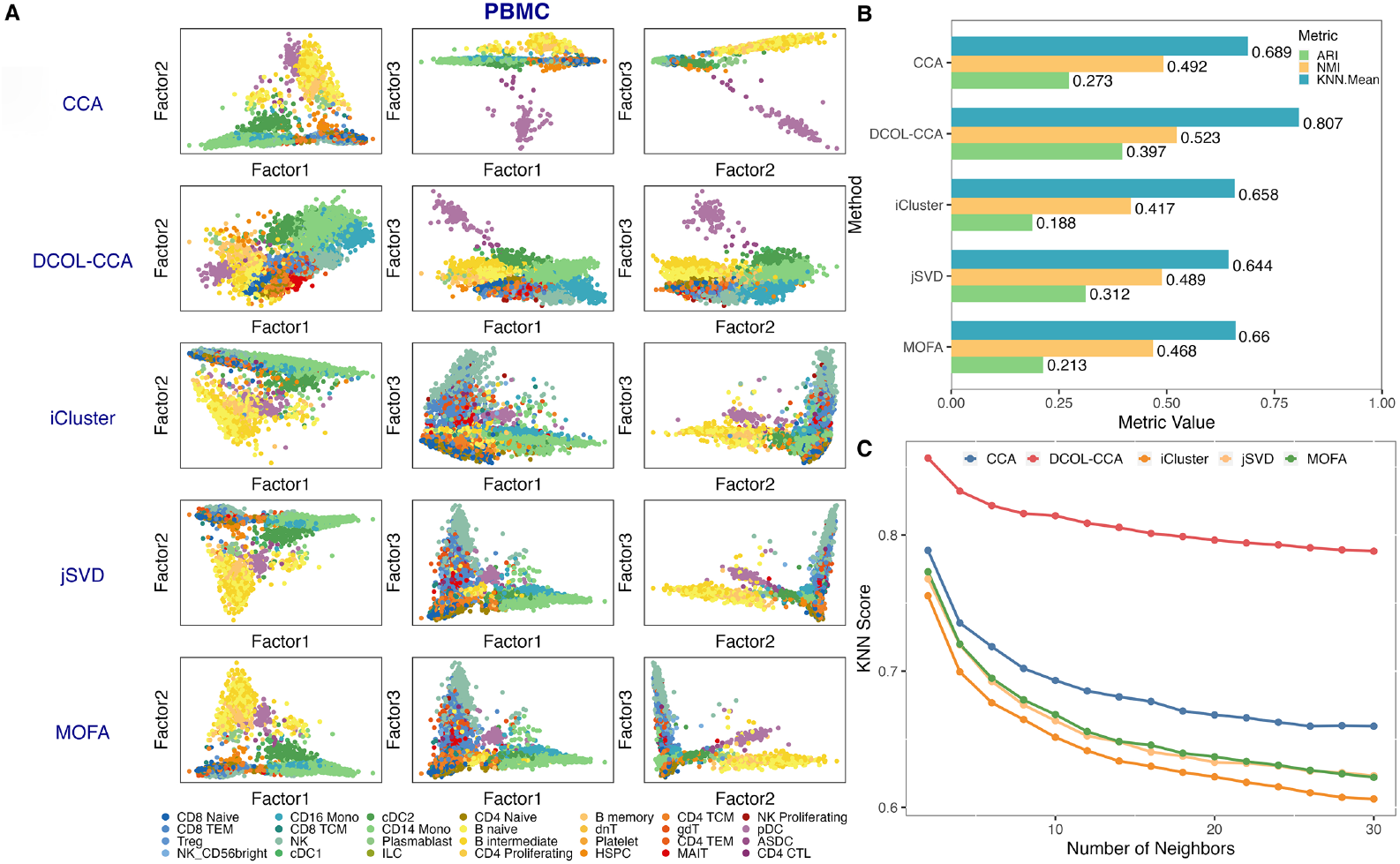
Joint dimensionality reduction results on PBMC dataset. (a) Pair-wise plots of the first three lower representations returned by each method. Points are colored based on cell types. (b) Clustering-based metrics ARI and NMI, as well as our new metric termed the KNN score, which calculates the proportion of nearest neighbors of each data point from the same cell type in the embedded data, were used to assess the performance of each DR method. (c) Comparison of KNN scores for different methods, with the number of neighbors (k) varying from 2 to 30.

The embedding plots from each method reveal that, except for specific cell types such as pDC cells, most cells from different groups tend to aggregate closely together without forming distinct cluster boundaries. Consequently, clustering-based metrics like ARI and NMI may underestimate the performance of the models. In light of this observation, we introduced a new metric termed KNN score to quantify and compare the performance of the methods. Specifically, we identified the k-nearest neighbors of each data point and calculated the mean proportion of neighbors belonging to the same cell group. By varying the number of neighbors from 2 to 30, we obtained the results shown in Figure 6C. Remarkably, DCOL-CCA consistently achieved significantly higher KNN scores compared to the other methods across all values of k, followed by CCA. The ARI and NMI scores (Figure 6B) also imply that DCOL-PCA leads the pack in performance. As illustrated by Figures 6A and 6C, jSVD and MOFA+ demonstrated very similar performances.

## Conclusion

Our study underscores the critical role of dimension reduction (DR) and joint dimension reduction (jDR) techniques in elucidating the complex relationships within and between single-cell omics data. While Principal Component Analysis (PCA) remains a cornerstone in linear DR, its limitations become apparent in handling diverse cell types. Nonlinear techniques such as UMAP and t-SNE have emerged as robust alternatives, each with distinct strengths and considerations. For joint analysis of multi-omics data, classical jDR methods often assume linear associations across assays, which may oversimplify the interactions between different biological components. The introduction of DCOL-PCA and DCOL-CCA in our study addressed these challenges, offering a robust solution for dimensionality reduction and integration of single-cell omics data. DCOL-PCA and DCOL-CCA’s capabilities to detect both linear and nonlinear relationships, along with their adaptability to single-omics and paired-omics data, position them as promising tools for researchers aiming to gain deeper insights into cellular heterogeneity.

Currently, DCOL-CCA is suitable for integrating paired-omics data, but it can be modified to analyze a larger number of omics data simultaneously. A natural extension is to maximize the sum of pairwise DCOL correlations to obtain lower-dimensional representations that account for the intricate interactions between different biological layers. This expansion would further enhance DCOL-CCA’s utility in deciphering complex cellular landscapes and advancing our understanding of cellular biology.

## Acknowledgments

This work was partially supported by the National Key R&D Program of China (2022ZD0116004), Guangdong Talent Program (2021CX02Y145), Guangdong Provincial Key Laboratory of Big Data Computing, and Shenzhen Key Laboratory of Cross-Modal Cognitive Computing (ZDSYS20230626091302006).

